# Stability of End-to-End Base Stacking Interactions in Highly Concentrated DNA Solutions

**DOI:** 10.1101/2023.01.25.525591

**Authors:** Sineth G. Kodikara, Prabesh Gyawali, James T. Gleeson, Antal Jakli, Samuel Sprunt, Hamza Balci

## Abstract

Positionally ordered bilayer liquid crystalline nanostructures formed by gapped DNA (GDNA) constructs provide a practical window into DNA-DNA interactions at physiologically relevant DNA concentrations; concentrations several orders of magnitude greater than those in commonly used biophysical assays. The bilayer structure of these states of matter is stabilized by end-to-end base stacking interactions; moreover, such interactions also promote in-plane positional ordering of duplexes that are separated from each other by less than twice the duplex diameter. The end-to-end stacked, as well as in plane ordered duplexes exhibit distinct signatures when studied via small angle x-ray scattering (SAXS). This enables analysis of the thermal stability of both the end-to-end and side-by-side interactions. We performed synchrotron SAXS experiments over a temperature range of 5-65 °C on GDNA constructs that differ only by the terminal base-pairs at the blunt duplex ends, resulting in identical side-by-side interactions while end-to-end base stacking interactions are varied. Our key finding is that bilayers formed by constructs with GC termination transition into the monolayer state at temperatures as much as 30 °C higher than for those with AT termination, while mixed (AT/GC) terminations have intermediate stability. By modeling the bilayer melting in terms of a temperature-dependent reduction in the average fraction of end-to-end paired duplexes, we estimate the stacking free energies in DNA solutions of physiologically relevant concentrations. The free-energies thereby determined are generally smaller than those reported in single molecule studies, which might reflect the elevated DNA concentrations in our studies.

**TOC Graphic:** 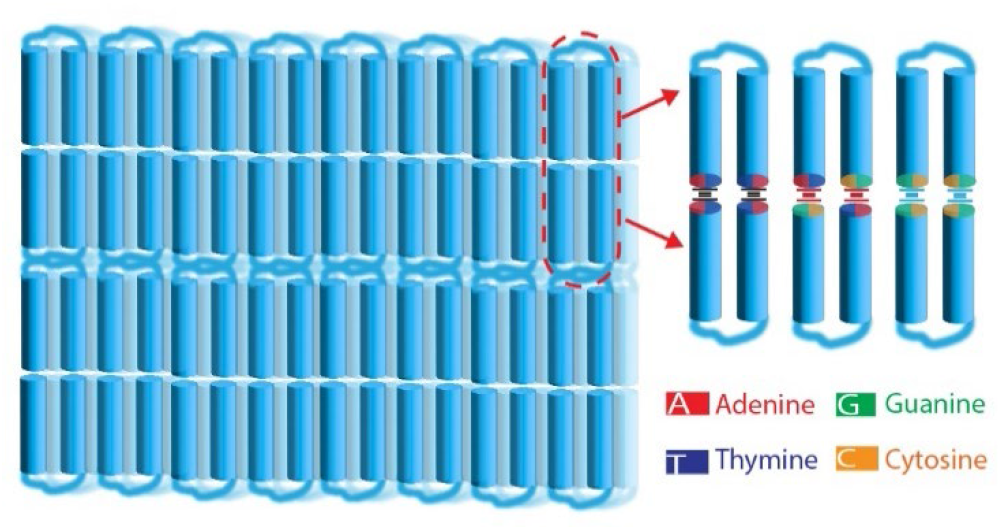

## INTRODUCTION

The local DNA concentration in chromosomes, viral capsids, and sperm nuclei is on the order of hundreds of mg/ml,^1^ concentrations at which DNA can form highly ordered lyotropic liquid crystalline (LC) phases. *In vitro*, short double-stranded DNA (dsDNA) fragments (∼6-20 bp, significantly shorter than persistence length of dsDNA) also exhibit LC phases, including chiral nematic and columnar mesophases.^2–8^

The introduction of a flexible single-stranded DNA spacer (“gap”) between two rod-like duplexes, producing a “gapped” DNA (GDNA) construct, has been the key for observing elementary layered (smectic) phases in concentrated DNA solutions.^9,10^ In addition to the DNA concentration (cDNA), the layer structure is sensitive to gap length and temperature.^9–11^ When the gap length is ≥ 10 nucleotides (nt), a bilayer structure composed of stacked duplexes prevails at ambient temperatures. The spacing between the bilayers slightly exceeds two duplex lengths, suggesting that attractive enthalpic interactions between blunt duplex ends and a net conformational entropy gain from segregating flexible “gap” segments act together to stabilize the bilayer stacking. At higher temperature, or for shorter gaps (∼3-9 nt), a monolayer smectic phase (approximately single duplex layer spacing) is observed. No smectic layer structure is observed in concentrated solutions when the gap length is ≤ 2 nt.^10^

For constructs with 10 nt or longer gaps, the positional order of the duplexes within the bilayers is short range (corresponding to a smectic-A LC phase) for DNA concentration in the range 230-260 mg/ml. At higher concentration (260-290 mg/ml) and at ambient temperature, the duplexes pack laterally in a hexagonal array within the bilayers (a smectic-B phase).^11^ Thus, at sufficient concentration, gapped constructs exhibit lateral positional order similar to the columnar phase formed by their fully paired analogs (FDNA), while also exhibiting the periodic layer structure characteristic of a smectic (unlike FDNA). The side-to-side interactions between duplexes that presumably stabilize the lateral order strongly depend on ionic content of the solution, a natural consequence of the high negative charge density of the DNA backbone, and are particularly sensitive to the presence of multivalent cations.^12,13^ Enhanced computational power and recent advanced in computational modeling of DNA and its interactions with cations and other DNA molecules have enabled realistic all-atom molecular dynamic simulations that probe such interactions at high effective DNA concentrations.^14–16^

The base stacking interactions that help stabilize the bilayer smectic phases in GDNA solutions are of fundamental importance for physiological pathways involving replication, repair, or restructuring of dsDNA. This has motivated measuring the strength of these interactions using different approaches.^17^ However, due to experimental limitations, small energies, and the difficulty in disentangling base pairing and base stacking interactions, this fundamental question is still actively investigated. To measure the strengths of these interactions various measurements have been performed, including gel electrophoresis assays on nicked dsDNA, thermal melting UV-Vis spectroscopy measurements on dsDNA constructs with different terminal overhangs, optical tweezers assays on DNA nanobeams with blunt duplex ends (where binding/dissociation of these ends is studied at different applied force), and centrifuge force microscopy assays on stacked DNA constructs (where dissociation of stacked constructs from the surface at different applied centrifugation speeds is monitored).^18–22^ The hydrophobicity of the DNA bases and hydrophilicity of the sugar-phosphate backbone are considered to be significant in deriving the stacking interactions between duplexes with blunt ends by creating hydrophobic pockets around the bases, which then attract each other when in close proximity (<2 nm).^23,24^ In the current study, we correlate the strength of these interactions with the stability of the bilayer structure in smectic GDNA solutions, which presents the opportunity to measure them in physiologically relevant DNA concentrations.

Fig. 1A shows the three GDNA constructs used in our study. Each consists of 48 bp duplex arms linked by a 20-thymine single-stranded “gap” segment. The constructs differ only by the terminal base pairs on the blunt ends of the duplex arms: AT or GC on both ends, or AT on one end and GC on the other. We shall refer to these constructs as AT-AT, GC-GC, and AT-GC, respectively. We performed SAXS measurements on samples with ∼280 mg/ml DNA concentration on beamline 11-BM at the National Synchrotron Light Source II, with the temperature varied from 5 to 65 °C in ∼5 °C steps. The range of scattering wavenumber (*q*) recorded in the SAXS measurements enabled monitoring the smectic layering (diffraction peaks at *q* of order the inverse duplex length) simultaneously with the lateral, in-layer order of the duplexes (*q* of order the inverse duplex diameter). We determine characteristic melting temperatures at which the bilayer structure and the in-layer ordering of the constructs in the smectic-B phase are destabilized. These results establish the central role of attractive end-to-end interactions between duplexes in stabilizing the bilayer phase. We also quantify the enthalpic and entropic contributions to the free energy associated with the bilayer ordering.

**Figure 1:**
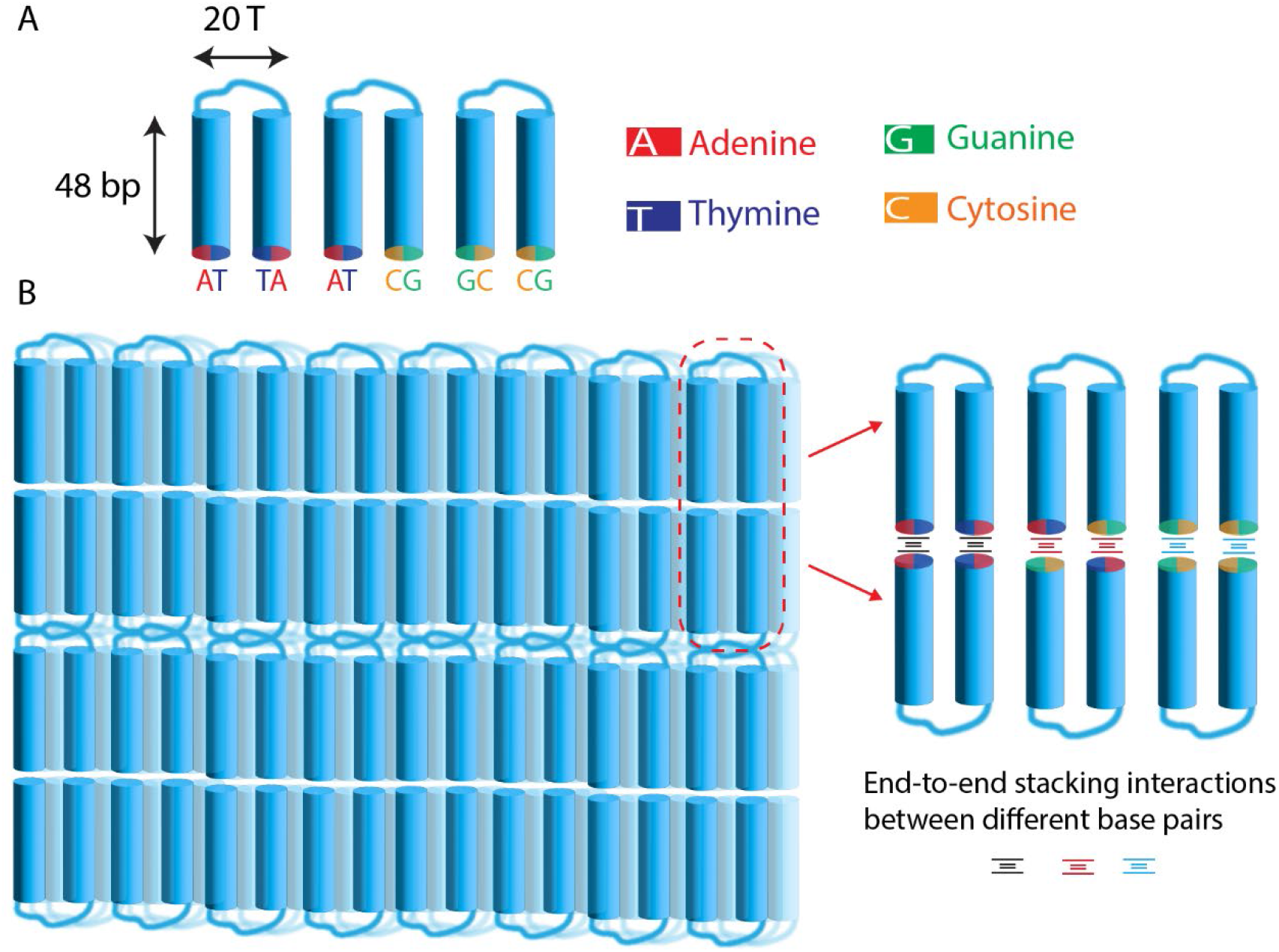
Schematic motifs of 48-20T-48 GDNA constructs, which have 48 bp long duplex arms connected with a flexible 20T long linker (“gap”). All constructs are identical except the terminal base pairs of the duplex arms. The constructs are labeled AT-AT, AT-GC, and GC-GC based on their terminal base pairs, which are color coded. (B) Schematics showing possible end-to-end stacking interactions for folded conformation of the GDNA constructs.

## RESULTS AND DISCUSSION

A typical SAXS pattern from our GDNA samples at ambient temperature features several sharp, small-angle peaks that represent multiple orders of diffraction due to the stacking of duplexes in smectic layers, along with a sharp peak at a wider angle that results from lateral positional ordering of the duplexes. In Fig. 2, azimuthally averaged SAXS intensity (on a log scale) vs *q* are plotted at various temperatures for the studied GDNA constructs. Four orders of small-angle diffractions are clearly detected. The wave number of the fundamental order (*q*_1_) corresponds to spatial periodicity 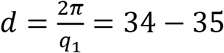 nm at 7 °C. The values of *d* are slightly greater than the 32.6 nm length of two 48-bp duplexes, and are consistent with a bilayer smectic structure.^10,11^ In addition, a sharp peak is recorded at wider angle. The position of this peak is consistent with the first order diffraction (wavenumber *q*_*w*_) from hexagonal lateral packing of the duplexes in the smectic-B phase of GDNA constructs previously reported at similar concentration and temperature.^11^ The corresponding center to center duplex spacings are 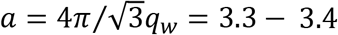 nm at 7 °C.

**Figure 2:**
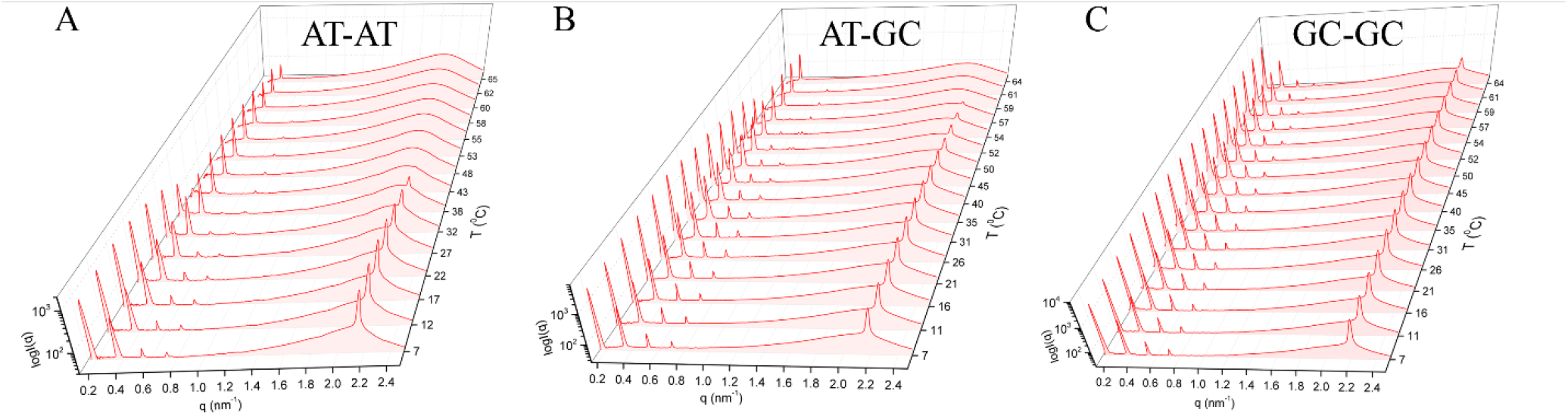
Temperature dependence of the azimuthally averaged SAXS intensity vs. scattering wavenumber *q* for GDNA samples. The intensity is on a log scale, and the data were acquired on a heating cycle (gradually increasing temperatures). (A) AT-AT construct; (B) AT-GC construct; and (C) GC-GC construct. All three samples are in a bilayer smectic-B phase at low temperatures, which melts at elevated temperature (indicated by diminishment of small-angle peaks at q_1_ ≈ 0.18 nm^-1^ and q_3_ ≈ 0.54 nm^-1^ that are specifically associated with the bilayer spacing and by the disappearance of the sharp wide-angle peak at q_w_ ≈ 2.2 nm^-1^, which is associated with in-layer positional ordering of the duplexes). While the bilayer peaks disappear at about 43 °C and 57 °C for AT-AT and AT-GC constructs, respectively, they do not completely melt for the GC-GC construct (at least up to 65 °C). The bilayer smectic-B melts into a monolayer smectic-A phase with characteristic small-angle peaks at q_2_ ≈ 0.36 nm^-1^ and q_4_ ≈ 0.73 nm^-1^, and a broad wide-angle peak indicating liquid-like short-range correlations within the layers.

Fig. 2A shows temperature-dependent intensity profiles vs *q* of x-rays diffracted at different angles for the AT-AT GDNA construct. The odd order (first and third order) peaks at *q*_1_ and *q*_3_ ≈ 3*q*_1_ completely melt along with the wider-angle peak (*q*_*w*_) at ∼43 °C while the even order peaks (second and fourth order peaks at *q*_2_ and *q*_4_ ≈ 2*q*_2_) survive up to (at least) 65 °C, the highest temperature studied, with a slight upward shift. The peaks at *q*_1_, *q*_3_, and *q*_*w*_ are therefore associated purely with the bilayer smectic-B phase. Their disappearance at ∼43 °C indicates a transition to a “monolayer” smectic-A phase, with layer spacing slightly greater than a single duplex and liquid-like correlations within the layers (accounting for the broad peak centered around a *q* value slightly below *q*_*w*_).

The intensity profiles for the AT-GC and GC-GC samples (Figs. 2B and 2C) evolve similarly with increasing temperature; however, the smectic-B to A transition temperatures are significantly higher than for the AT-AT sample. The bilayer peaks and wider-angle peak in the AT-GC sample are detectable up to ∼57 °C, while in the GC-GC sample these peaks (and hence the smectic-B layer structure) survive (at least) up to 65 °C. Evidently, the thermal stability of the bilayer smectic B phase is lowest for AT-AT termination on the duplex ends, highest for GC-GC, and intermediate between the two for AT-GC. This difference in bulk behavior among the three systems is due solely to the variation of single base pairs terminating the blunt ends of the duplexes, since otherwise the three GDNA constructs are identical, and the solutions were prepared in the same manner with consistent final component concentrations.

We analyzed the data in Fig. 2 by calculating the sum of the areas under the first and third order small angle peaks (*q*_1_ and *q*_3_) that are specifically associated with the bilayer stacking. As depicted in Fig. 3A, we first fit the data in each peak to a Gaussian function of *q*, and then integrated the fit result over *q* (excluding any contribution from the background scattering). Fig. 3B shows the square root of the resulting total integrated intensity under the peaks at *q*_1_ and *q*_3_, normalized to its maximum value and denoted 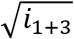, as a function of temperature for the three constructs studied. As argued in the Materials and Methods section, the normalized quantity 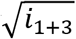 represents the average fraction (*f*) of end-to-end “bound” pairs of duplexes making up the GDNA bilayers. The argument assumes that the temperature-dependence of *f* is proportional to that of the amplitudes *ρ*_*n*_ contributing to the density wave, 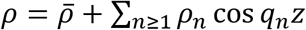 (*z* = axis normal to the layers, 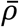 = average density), that describes the bilayer stacking. This assumption is reasonable if attractive end-to-end interaction between duplexes is the essential mechanism driving the bilayer formation. Only the peaks at *q*_1_, *q*_3_, which are associated with *ρ*_1_, *ρ*_3_, are included in the analysis, because at elevated temperature the peaks at *q*_2_, *q*_4_ are partially overlapped by diffraction (at slightly higher *q*) from the developing domains of the monolayer phase.

**Figure 3:**
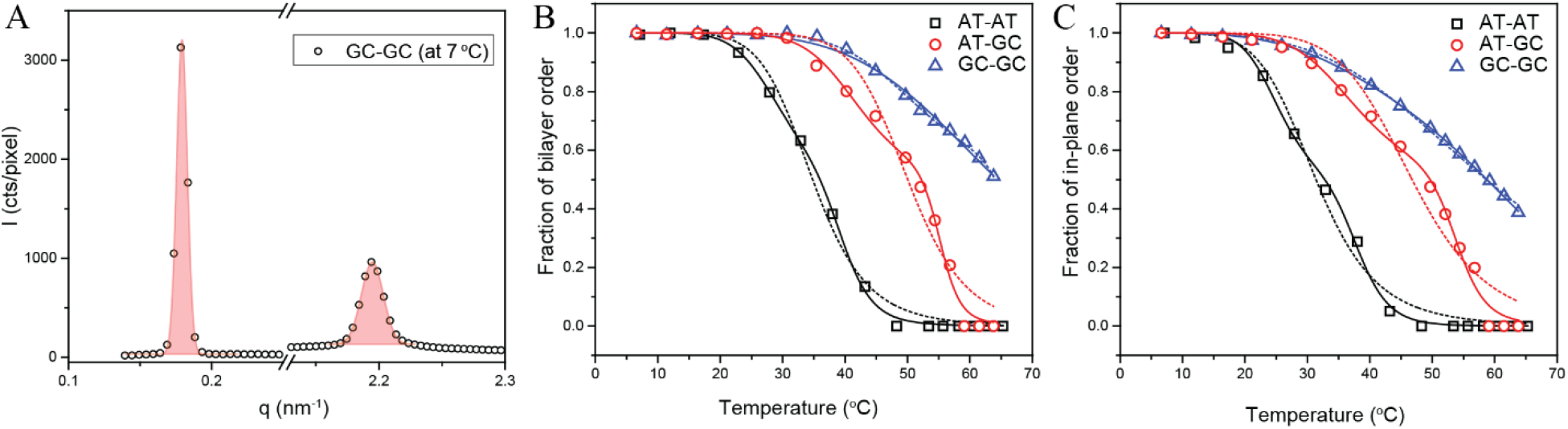
Analysis of the SAXS data in Fig. 2 as a function of temperature. (A) To determine the average fraction (*f*) of end-end “bound” duplexes constituting the bilayer domains, the areas under Gaussian fits to the small angle peaks at *q*_*1*_ and *q*_*3*_ (specifically associated with the bilayer structure) were calculated and summed. The results were normalized to their maximum value, yielding the quantity denoted *i*_1+3_ (see text). A similar analysis was performed for the sharp wide-angle peak at *q*_*w*_ (associated with in-layer ordering of the duplexes), yielding the quantity denoted *i*_*w*_ in the text. Two representative Gaussian fits are shown for the GC-GC construct at 7 °C for the peaks at *q*_*1*_ (at left) and *q*_*w*_ (at right). Open symbols are data points and solid lines are the Gaussian fits. The area under each peak is shaded. The base of the Gaussian represents the background. (B) Results for 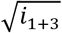 plotted vs temperature; as argued in the Materials and Methods section, we may associate this quantity with the fraction *f*. The dashed and solid lines are fits of the data points to a single and double Hill functions, respectively, from which characteristic melting temperatures *T*_*m*_ or *T*_*m*1_ and *T*_*m*2_ are determined. (C) Results for 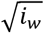 vs temperature. The data points were also fit to single and double Hill functions (dashed and solid lines, respectively) in order to obtain characteristic melting temperatures for comparison with those determined from 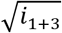.

For the three systems studied, Fig. 3B reveals that the fraction *f* decreases within different temperature ranges that correlate with the specific combination of base pairs terminating the duplex segments of a GDNA construct. Notably in the case of the AT-GC construct, the decrease appears to occur in two steps separated by an inflection point (“kink”) around 45 °C. The dashed and solid lines are fits of the data in Fig. 3B for 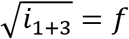 to single and double Hill functions,^25–27^ respectively. Expressions for the Hill functions are given in the Materials and Methods section; they are conventionally used to model the fraction of (non-covalently) bound pairs of interacting macromolecules in a solution as a function of temperature. Table 1 gives results for the fit parameter *T*_*m*_ (temperature at which half of the blunt duplex ends are stacked on average) from the fits using a single Hill function, and for the two parameters *T*_*m*1_ and *T*_*m*2_ determined from fits using a double Hill function. In all three systems studied, the double Hill function clearly provides a better description of the data than the single function, suggesting that the bilayers, formed by end-to-end paired constructs, melt via a “multi-step” process.

**Table 1.** Results for the fit parameter *T*_*m*_ from the fits using a single Hill function, and for the two parameters *T*_*m*1_ and *T*_*m*2_ determined from fits using a double Hill function.

|  | Bilayer Order Melting |  |  | In-plane Order Melting |  |  |
| --- | --- | --- | --- | --- | --- | --- |
|  | Single Hill<br>Function Fit | Double Hill<br>Function Fit |  | Single Hill<br>Function Fit | Double Hill<br>Function Fit |  |
| | $T_m(^{\circ}\text{C})$ | $T_{m1}(^{\circ}\text{C})$ | $T_{m2}(^{\circ}\text{C})$ | $T_m(^{\circ}\text{C})$ | $T_{m1}(^{\circ}\text{C})$ | $T_{m2}(^{\circ}\text{C})$ |
| **AT-AT** | 34.8±0.4 | 29.4±0.3 | 39.6±0.3 | 31.4±0.4 | 25.3±0.4 | 38.1±0.4 |
| **AT-GC** | 50.0±0.7 | 42.1±0.7 | 55.3±0.4 | 46.8±0.8 | 37.6±0.8 | 53.9±0.5 |
| **GC-GC** | 64.4±0.4 | 51.7±0.2 | 67.3±0.8 | 58.6±0.2 | 47.3±0.8 | 65.6±0.6 |

We can correlate the values for *T*_*m*_ determined from the Hill function analysis of 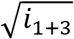 with values of the end-to-end interaction (“stacking”) energies, since higher stacking energies result in higher *T*_*m*_. The various stacking energies (see Materials and Methods section for nomenclature used to describe these energies) have been measured recently by single-molecule techniques.^18,22,28^ Using the values reported by Kilchherr *et al*.^18^ to compute an average stacking energy for each GDNA system studied, we find |*E*_*av*,AT-AT_ = −1.57| < |*E*_*av*,AT-GC_ = −1.94| < |*E*_*av*,GC-GC_ = −2.37| (in units of kcal/mol), which is exactly in line with the sequence of increasing melting temperatures *T*_*m*_ (Table 1) obtained from the Hill function fits to the data in Fig. 3. Here, *E*_*av*,XY-X’Y’_ represents the average energy of all stacking conformations that are relevant for our respective constructs. On the basis of this comparison, the key result from the single-molecule studies that GC-GC end-to-end interactions are on average significantly stronger than AT-AT interactions may now be extended to bulk systems of highly concentrated duplexes.

To discuss single versus multi step melting of the GDNA bilayers (and to help visualize the possible duplex stackings described above), it is useful to consider a specific model of a GDNA bilayer. We previously proposed^9–11^ that provided the gap segment is sufficiently long, the bilayers consist of folded GDNA constructs with the blunt ends of one construct paired with those of another (as illustrated in Fig. 1B). Figs. 4A-C show schematically six different ways our GDNA constructs can fold and pair. These can be divided into two groups based on the strength of the associated stacking interactions. Fig. 4A illustrates the groups for the AT-AT construct. The purple (cyan) shading distinguishes duplexes with the terminal A in the 5′ (3′) position. In Group 1, the stacking energies for each pair of opposing duplexes are degenerate and equal to *E*_*TA*:*AT*_. In Group 2, the energies are either *E*_*TA*:*AT*_ (purple duplexes stacked against each other) or *E*_*TA*:*AT*_ (cyan stacked against cyan). As indicated in Figs. 4B and C, stacking interactions among paired, folded AT-GC and GC-GC constructs may be similarly grouped.

**Figure 4:**
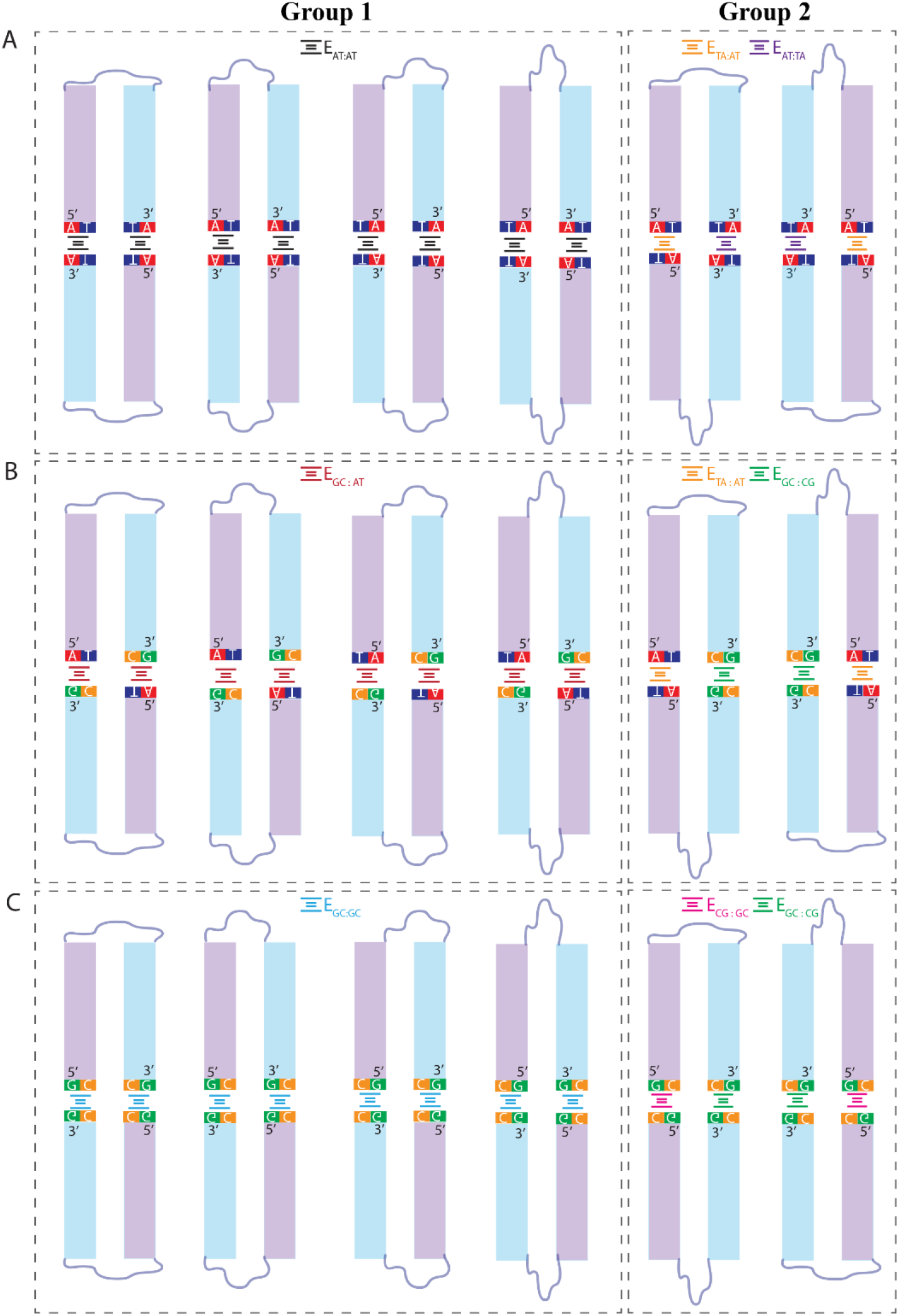
Schematics showing possible end-to-end stacking interactions for folded conformations of the AT-AT construct (A); for AT-GC construct (B); and for GC-GC construct (C). The duplex arms are color coded such that duplexes that have an A or a G at 5′ (3′) positions are purple (cyan). The conformations of each construct are classified as Group 1 and Group 2. The three conformations in Group 1 have degenerate stacking energies, while the two conformations in Group 2 have distinct stacking energies, resulting in a total of three distinct stacking energies for each construct.

Again using the results of single-molecule measurements in Kilchherr et al^18^, the average stacking energy in a pair of folded AT-AT constructs is 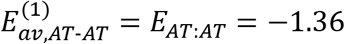 kcal/mole for Group 1 (indicated by the superscript 1) and 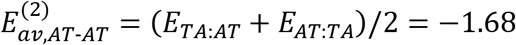 kcal/mol for Group 2. Similarly, we get 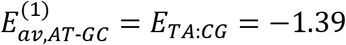 kcal/mol, 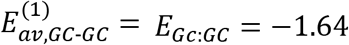 kcal/mol and 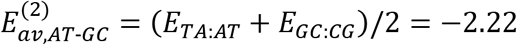 kcal/mol, 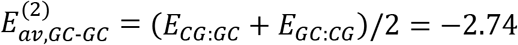 kcal/mol for Groups 1 and 2 of the AT-GC and GC-GC constructs. We can then argue that the bilayers in each case melt in two steps characterized by temperatures *T*_*m*1_, associated with melting of domains in Group 1 (lower stacking energy cost), and *T*_*m*2_, associated with melting of Group 2 domains (higher stacking energy cost). As discussed above and evident in Fig. 3B, the corresponding analysis of 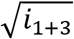 with two Hill functions describes the data well. Moreover, the ascending order of values of *T*_*m*1_ and *T*_*m*2_ match the ordering of 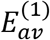 and 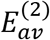, respectively, among the three systems studied. However, the argument (and particularly the definition of 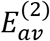) assumes that the discrete pairs of constructs in the two groups shown in Figs. 4A-C have an effective end-end interaction that is a hybrid of the component end-end stackings. If, on the other hand, the bilayer structure is formed from stacking extended (“unfolded”) constructs, or if the folded pairs are not discrete (*e*.*g*., single ends of one folded construct paired with single ends of two others)^10^, this assumption is questionable, and the bilayer melting may instead be determined by some other combination(s) of the non-degenerate end-end interaction energies. In any case, given their limited resolution, the present data are adequately described by an analysis based on a two-step model.

Based upon end-to-end unbinding of stacked duplexes as the mechanism for the melting of the bilayer smectic, we can calculate the Gibbs free energy of stacking interactions. As described in the Materials and Methods section, we first obtain the effective equilibrium “binding” constant characterizing the duplex end-end stacking interactions, 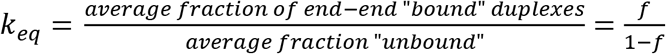, as a function of temperature, using the data for *f* in Fig. 3. Fig. 5 shows the results for In *k*_*eq*_ plotted against 1/*T*. Fits of these results to the van’t Hoff equation^29,30^, ln 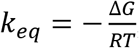, yield the following Gibbs free energies evaluated at 296 K: Δ*G* = −0.45 ± 0.03 kcal/mol for the AT-AT construct, −1.54 ± 0.16 kcal/mol for the AT-GC construct, and −1.82 ± 0.16 kcal/mol for the GC-GC construct. The entropic and enthalpic contributions to Δ*G* at 296 K are listed in Table 2. These values reinforce the conclusion that the GC-GC stacking interactions are the most stable, while AT-AT are the least, with AT-GC intermediate between the two.

**Table 2.** Results of the analysis of data presented in Fig. 5. The free energies are calculated at 296 K (23 °C).

| **Construct** | **Intercept**<br>**(kCal/mol)** | **Slope (K)** | **$\Delta H$**<br>**(kCal/mol)** | **$\Delta S$**<br>**(kCal/mol/K)** | **$\Delta G$**<br>**(kCal/mol)** |
| --- | --- | --- | --- | --- | --- |
| **AT-AT** | 65.30±3.04 | -19556±913 | 38.84±1.81 | 0.13±0.01 | 0.45±0.03 |
| **AT-GC** | 49.53±3.57 | -15435±1130 | 30.65±2.24 | 0.10±0.01 | 1.54±0.16 |
| **GC-GC** | 38.45±2.38 | -12296±765 | 24.42±1.52 | 0.08±0.01 | 1.82±0.16 |

**Figure 5:**
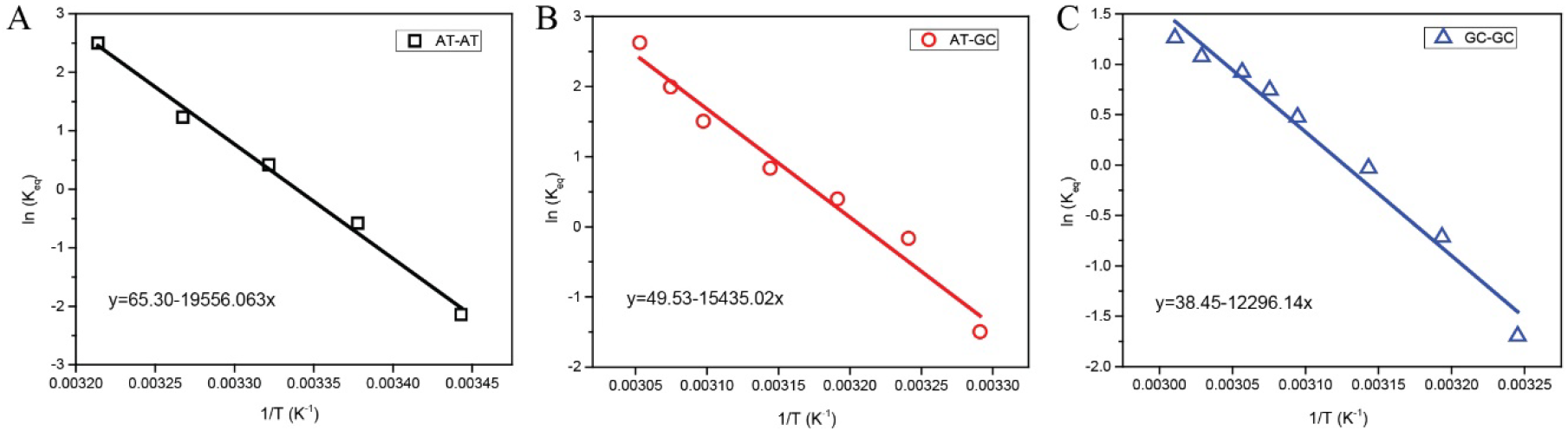
Calculation of Gibbs free energy due to end-to-end stacking of duplexes. Data and linear fit for AT-AT construct are shown in (A); for AT-GC construct in (B); and for GC-GC construct in (C). Symbols are results for the natural logarithm of effective equilibrium “binding” constant (*k*_*eq*_) obtained from the results for 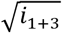 (see Materials and Methods section), and the solid lines are linear fits. The best-fit equation is given in each figure.

Free energy measurements using other approaches^18–21^ show that stacking of purines with purines is more stable than stacking of pyrimidines with pyrimidines and specifically that GC-GC stacking is more stable than AT-AT stacking, with AT-GC stacking being between these two limits, in agreement with our results. The variation in assay conditions, particularly ionic conditions, and level of decoupling between base-pairing and base-stacking interactions make it challenging to perform a more detailed quantitative comparison. To illustrate, 20 mM MgCl_2_ or 500 mM NaCl were used in one study,^18^ while 50 mM phosphate buffer saline (ionic strength ∼110 mM) was used in another,^22^ and 10 mM MgCl_2_ and a few mM KCl were used in yet another study.^28^ Using thermal melting studies, Yakovchuk *et al*. characterized variation of stacking free energy with monovalent cation concentration for different base configurations, and demonstrated that the free energies become ∼30% more negative (more stable stacking) as the salt concentration was increased from 15 to 100 mM NaCl.^21^ They also demonstrated that base-stacking and not base-pairing was primarily responsible for the enhanced stability at higher ionic strength. Furthermore, Kilchherr *et al*.^18^ compared the breaking time of their stacked nanobeams in 20 mM MgCl_2_ to that in 500 mM NaCl and observed that the stacking interactions were more stable in 20 mM MgCl_2_, confirming the effect of divalent cations in lowering Δ*G*. In general, the free energies we report are on the less negative end of the spectrum of values from the studies cited above, which is consistent with the absence of divalent cations in our samples.

Another significant parameter which would impact the stacking free energies is the level of crowding of the environment, which would particularly influence the entropic contributions to the free energy. Most of the recent studies we cite^18,22,28^ are single molecule studies where there is effectively zero interaction between neighboring stacked duplexes; however, even typical bulk studies use DNA concentrations that are ∼3 orders of magnitude lower than those we use (*e*.*g*. ∼ µM in thermal melting assays^21^ *vs*. mM in our studies, which is required for LC phase formation). Our measurements are unique in this respect as they probe base stacking interactions at physiologically relevant DNA concentrations. Relatedly, all free energies we report are based on transition from a bilayer smectic phase to a monolayer smectic phase, which in our model (Fig. 1B) results from breaking pairs of stacked duplex arms. Therefore, as described earlier, the free energies we report have inherent averaging, which is not the case in the single molecule approaches.

In order to determine characteristic melting temperatures of the in-layer order that can be compared to the *T*_*m*_ values associated with the bilayer melting, we calculated the area under the sharp, wider-angle peak (*q*_*w*_) in Fig. 2 in similar fashion to the area under the small angle peaks, as illustrated in Fig. 3A. Fig. 3C shows the results for the square-root of integrated intensity 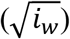 vs temperature, together with fits to single and double Hill functions. Values for the parameters *T*_*m*_ or *T*_*m*1_ and *T*_*m*2_ are listed in Table 1. These values are systematically lower than those for the bilayer peaks for all three GDNA constructs, which indicates that, in heating, the in-layer order starts to melt before end-to-end stacking interactions are broken. Similarly, when cooling down, the duplex ends stack before in-layer order emerges. These findings imply end-to-end stacking interactions stabilize first and promote the lateral ordering of duplexes during formation of the bilayer smectic-B phase.

It should be noted that the measurements reported in this work were performed in the absence of any multivalent cations, which are proposed to enhance attractive interactions between duplexes when present at sufficiently high concentrations. Even though the enhancement is expected to be negligible at physiologically relevant ion concentrations,^31^ it would be of fundamental interest to investigate the impact of multivalent cations on the stability of both bilayer and lateral ordering of GDNA duplexes.

## CONCLUSIONS

Based on SAXS measurements on concentrated solutions of gapped DNA constructs differing only in terminal base pairs at the blunt ends, we showed that the thermal stability of the bilayer smectic-B phase in GDNA depends on the strength of the duplex end-to-end stacking interactions. While not systematically investigated in this study, the reduced DNA terminal breathing in a GC base pair, compared to an AT base pair, likely impacts the stability of stacking interactions and that of the bilayer smectic-B phase as well. Lateral order of the duplexes consistently onsets at a lower temperature than their end-to-end stacking. GC-GC terminated constructs produce the most stable bilayers, while the AT-AT constructs yield the least stable. AT-GC constructs form bilayers with intermediate stability. We also presented evidence of a multi-step bilayer melting process, which we rationalized using a model of bilayers composed of folded, end-to-end paired GDNA constructs – a model previously introduced to describe elementary smectic ordering of GDNA with sufficiently long gap segments.

The present study demonstrates that the relative strengths of base stacking interactions can be studied at physiologically relevant DNA concentration, temperature range, and ionic conditions via the stability of smectic LC phases formed by gapped DNA constructs. In addition to their significance in liquid crystalline phase formation, base stacking interactions are of fundamental biological significance; our results provide important insights for modeling efforts involving these interactions. The elevated stability of the smectic phases, particularly those formed by stacking GC-GC terminated duplexes, also suggests potential use in biosensor applications where physiological temperature range is of primary focus.

## MATERIALS AND METHODS

### GDNA Synthesis

This study utilized three different GDNA constructs, all consisting of 48-base pair duplexes bridged by a single-strand gap of 20 Thymine bases (48-20T-48) with either AT or GC base pairs terminating each of the duplex arms (Figs. 1A-C). The GDNA constructs were formed by annealing three PAGE-purified strands (O1, O2, and O3), purchased from ExonanoRNA (Columbus, OH) or GenScript (Piscataway, NJ). The sequences of O1, O2, and O3 are as follows:

**O1 Strand**: 5′-**<u>N</u>**CAGATGCACATATCGAGGTGGACATCACTTACGCTGAGTACTTCGAA

TTTTTTTTTTTTTTTTTTTTAAGCTTCATGAGTCGCATTCACTACAGGTGGAGCTATAC

ACGTAGAC**<u>M</u>**

**O2 Strand**: 5′-TTCGAAGTACTCAGCGTAAGTGATGTCCACCTCGATATGTGCATCTG**<u>K</u>**

**O3 Strand**: 5′-**<u>R</u>**GTCTACGTGTATAGCTCCACCTGTAGTGAATGCGACTCATGAAGCTT

For AT-AT Construct: **<u>N</u>**=A, **<u>M</u>**=A, **<u>K</u>**=T, and **<u>R</u>**=T

For AT-GC Construct: **<u>N</u>**=A, **<u>M</u>**=G, **<u>K</u>**=T, and **<u>R</u>**=C

For AT-GC Construct: **<u>N</u>**=G, **<u>M</u>**=G, **<u>K</u>**=C, and **<u>R</u>**=C

The GDNA synthesis process, sample loading procedure into the borosilicate X-ray capillaries, and details of the SAXS measurements are described in Supplementary Information and in previous publications^10,11^.

### Nomenclature for Stacking Energies

The “AT-AT” construct in Fig. 1A has AT as the terminal base pair on both duplex arms, which results in AT-AT type stacking between neighboring GDNA constructs. The “AT-GC” construct (Fig. 1A) has an AT terminal base pair on one of the duplexes and a GC base pair on the other duplex. This results in a mix of stacking interactions where AT base pairs could stack with either AT or GC base pairs, and vice versa. The “GC-GC” construct (Fig. 1A) has GC as the terminal base pair on both duplex arms, which results in GC-GC type stacking.

In the AT-AT system, there are three distinct stacking energies. One, designated *E*_*TA*:*AT*_, corresponds to A’s (T’s) lining up against A’s (T’s), with the 5′ (3′) ends of the single strands on one duplex opposing the 3′ (5 ′) ends on the other. Here (and below) the subscript *X*: *X*′*X*′ denotes 3′-*X* stacked opposite 5′-*X*′ and 5′-*X* stacked opposite 3′-*X*′. Two additional stacking energies are associated with A’s (T’s) lining up against T’s (A’s): *E*_*TA*:*AT*_ or *E*_*AT*:*TA*_ if the terminal A’s are in the 5′ or 3′ positions, respectively. Analogously, the distinct stacking energies for the GC-GC system may be designated *E*_*GC*:*GC*_, *E*_*CG*:*GC*_, and *E*_*GC*:*CG*_. Finally, the AT-GC system introduces four additional non-degenerate stacking energies, *E*_*AT*: *CG*_, *E*_*AT*:*CG*_, *E*_*AT*:*GC*_, and *E*_*AT*: *CG*_. However, only the last of these (*E*_*AT*: *CG*_) is relevant in our samples; the others apply when the A and G bases are both in the 5′ or 3′ positions, or when A is in the 3′ and G in the 5′ position – none of which are the case for our construct. Also, in this nomenclature *E*_*XY*:*X*′*Y*′_ = *E*_*Y*′*X*′:*YX*_, *e*.*g*., *E*_*TA*:*CG*_ = *E*_*GC*:*AT*_, as they both describe stacking of the same duplexes where the bases of one or the other duplex are registered first. Therefore, not every possible combination is written explicitly in this description. In addition to *E*_*TA*:*CG*_, stacking interactions with energies *E*_*TA*:*AT*_ or *E*_*GC*:*CG*_ are possible in our AT-GC system.

### X-ray scattering intensity from bi- and mono-layer stacking of GDNA constructs

The mass density wave describing the layer structure may be written as 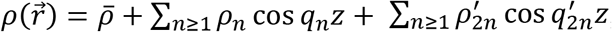, where 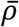 is the average density, *ρ*_*n*_ and 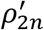 are real, non-negative Fourier amplitudes for bi- and mono-layer contributions, respectively, to the overall density, *z* is the direction of the layer normal, *q*_*n*_ = *nq*_0_, and 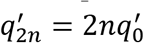 with 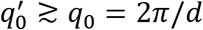 (*d* = bilayer periodicity). (The latter inequality reflects the slight upward shift in the peaks at 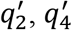 observed in the monolayer smectic-A phase compared to *q*_2_, *q*_4_ in the smectic-B phase.) The intensity of X-rays scattered from 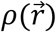 is 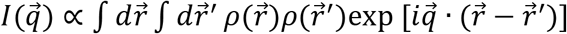. Inserting the expression for 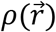 and taking 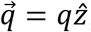, we obtain a series of delta functions in *q* − *q*_*n*_, which experimentally are broadened both by the finite size of smectic domains and by limited resolution in *q*. The finite size effect can be accounted for by replacing the delta function peaks with Gaussian lineshapes centered on each *q*_*n*_ and with widths that scale with the inverse square of the characteristic size *L* = *Nd* of the domain along the layering direction (Warren approximation, *N* = number of bilayers spaced by *d*).^32^ Convolution with a Gaussian resolution function (whose width is estimated from the experimental conditions to be about 1.5 times the spacing between data points in Fig. 2) broadens the lineshape, while maintaining a Gaussian profile. Fitting the peaks at *q*_1_ = *q*_0_ and *q*_3_ = 3*q*_0_ to Gaussian profiles and integrating the fitted profiles over *q* gives 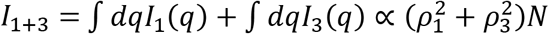. (A factor of *N*^2^ that comes from the Warren approximation is reduced to *N* after the integration over *q*.) Normalizing *I*_1+3_ to its maximum value at low temperature then yields the quantity *i*_1+3_ as a function of temperature.

### Characteristic Melting Temperature and Free Energy Calculations

The results for *i*_1+3_ were analyzed assuming that the Fourier amplitudes *ρ*_1_ and *ρ*_3_ (related specifically to bilayer stacking) are proportional the temperature-dependent, average fraction *f* of end-to-end paired (“bound”) duplexes in a typical sample volume, whence *I*_1+3_ *∝ f*^2^. The characteristic number *N* of bilayers in a finite sized domain could also vary with temperature (*T*), but we did not to observe any significant change with *T* of the SAXS peak widths at *q*_1_ and *q*_3_. (Within the experimental resolution in *q*, the peak widths should scale as *N*^−2^.) Normalization to the maximum *I*_1+3_ gives 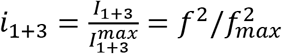, and then assuming *f*_*max*_ = 1 (at sufficiently low temperatures), we get *i*_1+3_ = *f*^2^. Values of 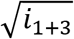 versus temperature *T* were fit to single or double Hill functions, which were used to model *f*(*T*) – specifically, 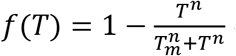 or 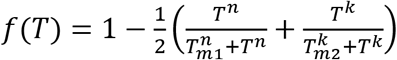, where *T*_*m*_, or *T*_*m*1_ and *T*_*m*2_, are characteristic melting temperatures, and *n* and *k* are positive exponents.

The free energies were calculated from the van’t Hoff equation.^29,30^ Briefly, the Gibbs free energy (Δ*G*) can be related to entropy (Δ*S*) and enthalpy (Δ*H*) as follows: Δ*G* = Δ*H* − *T*Δ*S*, where *T* is the absolute temperature. Δ*G* may also be expressed in terms of an equilibrium “binding” constant (*k*_*eq*_) as Δ*G* = −*RT* In *k*_*eq*_, where *R* = 1.987 cal/mol/K is the universal gas constant and 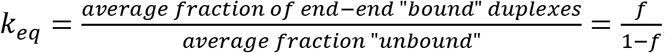. Therefore: In 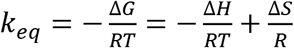. We calculated In *k*_*eq*_ as a function of *T* from the data for 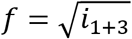 (Fig. 3) and then plotted In *k*_*eq*_ as a function of 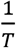 for temperatures in the transition region (Fig. 5). After fitting the result to a line and determining Δ*H* from the slope and Δ*S* from the intercept, we calculated Δ*G* for *T* = 296 K.

## Acknowledgements

The research reported here was supported by the National Science Foundation under grant DMR-1904167. The authors are particularly grateful to Ruipeng Li and Masa Fukuto for their assistance in performing the SAXS/WAXS measurements on the CMS beamline (11-BM) at the National Synchrotron Light Source II, a U.S. Department of Energy (DOE) Office of Science User Facility operated for the DOE Office of Science by Brookhaven National Laboratory under Contract No. DE-SC0012704.

## Supporting Information Available

Details of GDNA synthesis and SAXS measurements

