## Supporting Information for "Stability of End-to-End Base Stacking Interactions in Highly Concentrated DNA Solutions"

**DNA Sample Preparation**

Polyacrylamide gel electrophoresis (PAGE) purified oligomers (O1, O2, and O3 listed in manuscript) were obtained from commercial ExonanoRNA or GenScript. The three strands were annealed at equimolar concentrations (10 μM) at 90 °C for 10 min in a 150-mM NaCl, 10-mM trisaminomethane-HCl (pH 7.5), and 0.1-mM ethylenediaminetetraacetic-acid aqueous buffer. The mixture was then slowly cooled to room temperature over many hours (natural cooling rate of a ~3 L water bath).

The annealed sample was then passed through a 50-kDa membrane filter (Amicon Ultra from Millipore) by centrifuging at 5,000 rpm for 15 min. The supernatant, with a volume of ∼100 μL and salt concentration of 150 mM NaCl, was then collected. Next, we added deionized water to the supernatant solution, increasing the total volume by approximately five times and reducing the NaCl concentration to ∼30 mM NaCl. This solution was then concentrated by passing it through a 10-kDa filter (Amicon Ultra from Millipore) and centrifuging it for 12 min at 12,000 rpm, which maintains the NaCl concentration at ∼30 mM NaCl. Using a NanodropOne instrument (Thermo Scientific), we measured the GDNA concentration of the concentrated supernatant in the range 80−100 mg/mL. We made minor adjustments to salt and GDNA concentrations of this supernatant, so that ∼150 mM NaCl would be reached at a target DNA concentration at 280 mg/mL (after evaporation of water from the open end of the capillary).

Before loading the GDNA samples, we calibrated the capillaries by adding known volumes of deionized water and measuring their height in the capillary. The calibrated capillaries were dried, and the supernatant GDNA samples were gradually and carefully pipetted in.

Loaded capillaries were placed in a custom-made aluminum holder that was partially immersed in a water bath at 40 °C to 45 °C; both the GDNA and NaCl concentrations increase as the water evaporates, and the solution height decreases. When the desired height, and hence, desired GDNA concentration, was reached, the capillary was removed from the bath, and the open end was sealed with an inert epoxy. The sealed capillaries were kept at 4 °C until the X-ray measurements, and the sample height was checked periodically to confirm integrity of the sealing.

**SAXS Measurements**

SAXS measurements were performed on beamline 11-BM at the National Synchrotron Light Source II. The incident X-ray energy was 17 keV, and the incident beam size at the sample was 200 μm×200 μm. The typical acquisition time for SAXS patterns was 5 s and we did not observe any evidence of X-ray damage to samples. Commercial hot/cold stages with Kapton film windows were used to regulate the sample temperature between 5 °C and 65 °C. Background scattering was recorded from a capillary containing pure buffer solution, which was subtracted from the data taken on the GDNA samples. A capillary filled with silver behenate powder was used to calibrate the scattering wave number (q) in the plane of the detector.
